# The effect of feeding behavior of *Monochamus alternatus* (Coleoptera: Cerambycidae) on the departure of pine wood nematode, *Bursaphelenchus xylophilus* (Nematoda: Aphelenchoididae)

**DOI:** 10.1101/2019.12.13.875575

**Authors:** Yang Wang, Fengmao Chen, Lichao Wang, Lifeng Zhou, Juan Song

## Abstract

In order to study the causes of pine wood nematode (PWN) departure from *Monochamus alternatus*, the effects of the feeding behavior of *M. alternatus* on the start date of the departure of PWN were studied. The start date of the departure of PWN carried by the directly fed *M. alternatus* was 5—13 d after beetle emergence, mainly concentrated within 6—10 d, with a mean (±SD) of 8.02 ± 1.96 d. The start date of the departure of PWN carried by the *M. alternatus* fed after starvation was 5—14 d after beetle emergence, mainly concentrated within 6—9 d, with a mean of 7.76 ± 2.28 d. The results show that there was no significant difference in the start departure date of PWN between the two treatments. This shows that the feeding behavior of *M. alternatus* is not the trigger for PWN departure. At the same time, it was found that the motility of the PWN carried by *M. alternatus* at 8 d after emergence was significantly greater than that of the PWN carried by the newly emerged *M. alternatus.* And the PWN carried by *M. alternatus* at 8 d after emergence was extracted more easily than the PWN carried by newly emerged beetles. These results show that greater motility was associated with easier departure of PWN from *M. alternatus.* In addition, transcriptome sequencing found that the level of oxidative phosphorylation metabolism of PWN carried by beetles at 8 d after emergence was significantly higher than that in the PWN carried by newly emerged beetle. High oxidative phosphorylation was associated with increased energy production and motility by the PWN and were the internal cause of the start of nematode departure.

## Introduction

Pine Wilt Disease (PWD) constitutes one of the most serious conifer diseases worldwide, affecting *Pinus* spp. from the Far East forestlands (Japan, China and Korea) (Cheng et al., 1983; Yi et al., 1989), and from Europe (Portugal and Spain) (Abelleira et al., 2011; Fonseca et al., 2012; Mota et al., 1999; Robertson et al., 2011). This disease causes significant economic and environmental damage to the countries affected, with large annual losses of timber (Mamiya et al., 2004), increased cost in management procedures, including disease and pest control (Mamiya et al., 2004; Yang et al., 2004). This disease is caused by the pine wood nematode (PWN), *Bursaphelenchus xylophilus* (Steiner & Buhrer) Nickle, and PWN transmission is dependent on vector insects, of which the main vector insect in East Asia is *Monochamus alternatus* Hope (Linit, 1989; Morimoto and Iwasaki, 1972). There are two developmental forms in the life cycle of the juvenile PWN, namely the propagative and the dispersal forms. Under favorable conditions, PWN molt into their propagative form, which then reproduce rapidly (Mamiya, 1984; Wingfield, 1983). However, under unfavorable conditions, e.g., high population density, nutritive deficiency, desiccation, and low temperature, the propagative second-stage juveniles will molt to produce dispersal third-stage juveniles, which will aggregate around the pupal chamber of the vector insects (Linit, 1988; Zhao et al., 2014). The dispersive third-stage juveniles molt to produce fourth-stage juveniles, which enter the tracheal system of the vector as the adult beetles emerge (Linit, 1988,1990; Necibi and Linit, 1998; Zhao et al., 2014); when the adult beetles feed on healthy pine trees, the fourth-stage dispersive juveniles to be transmitted to the healthy pine trees through wounds caused by the vector (Niu et al., 2012; Stamps and Linit,1988,2001; Zhao et al., 2007).

The PWN cannot be transmitted immediately after the emergence of *M. alternatus.* A number of studies have investigated the time after *M. alternatus* emergence when PWN transmission starts. Fourth-stage dispersive juveniles exit the tracheal system of *M. alternatus* at 3–5 d (Hosoda and Kobayashi, 1977), at 5 d (Jikumaru and Togashi, 2000), at 7 d (Aikawa, 2008), at 7–12 d (Wang et al. 2019), at 10 d (Togashi, 1985), or within 10 d after the emergence of the vector beetles (Enda, 1972).

What triggers the departure of PWN from its insect host? A number of studies have investigated various factors: the degradation of neutral storage lipid was correlated with nematode exit (Stamps and Linit, 1988); assumed *β*-mvrcene to play an important role in the transmigration of the PWN from the sawyer to the pine tree (Ishkawa et al., 1986); PWN has a trait of spontaneous departure from *M. alternatus,* and pine volatiles repress PWN departure from *M. alternatus*(Aikawa and Togashi, 1998); PWN departure behavior possibly endogenous natural factors (Aikawa and Togashi, 1998; Stamps and Linit, 1988); when the CO_2_ concentration in the trachea of vector beetle reaches a certain critical value, PWN begins to escape from the beetle (Wu et al., 2019). The feeding period is an important stage in the life cycle of the vectors, as well as a key step in PWN transmission (Togashi and Shigesada, 2006; Yoshimura et al., 1999), although whether the feeding behavior of vector is the factor triggering PWN departure has not been studied. In order to further study the causes of PWN departure from vector, we investigated the effect of the feeding behavior of *M. alternatus* on PWN departure, the differences in motility and difficulty level of extraction of the PWN carried by *M. alternatus* at 8 d after emergence and newly emerged, and differences in the PWN transcriptome at these two time nodes.

## Materials and methods

### Collection of infected wood and *M. alternatus*

In March 2019, dead specimens of *Pinus massoniana* trees, infested by *M. alternatus* larvae and PWN, were collected at Bocun Forest Farm, Huangshan City, Anhui Province in eastern China. The pine trees were cut into logs (1.0—1.2 m in length) and maintained in outdoor insect cages (0.5 m × 0.5 m × 1.2 m), as shown in Fig. 1 A. *M. alternatus* adults were collected daily (every 6 h) during the period of adult beetle emergence from the logs.

**Fig. 1.**
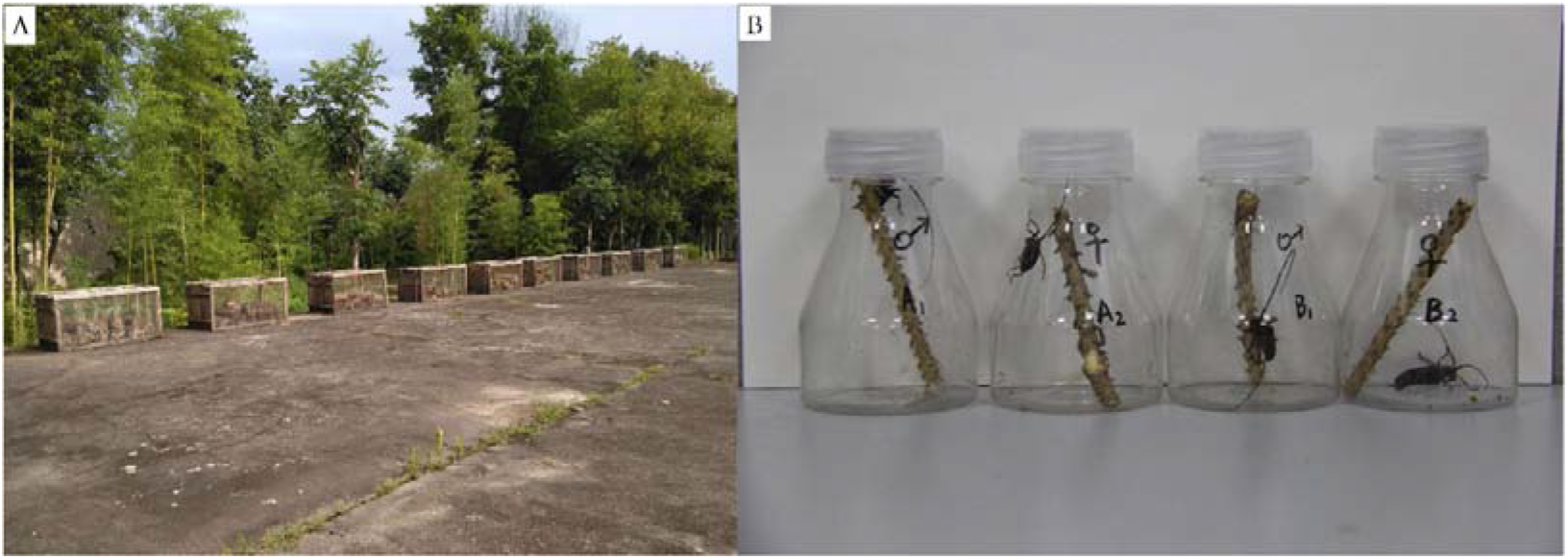
Collection and feeding of *Monochamus alternatus*. A: The dead pine trees were cut into logs and maintained in outdoor insect cages. *Monochamus alternatus* adults were collected daily (every 2 h) during the period of adult beetles emergence. B: The collected *M. alternatus* adults were divided into two groups and maintained in culture flasks at 25°C, one of which was fed (with fresh one-year-old pine twigs with the needles removed) immediately after collection, whereas the second group was fed after a starvation period of 4 d.

The collected *M. alternatus* were divided into two groups and maintained in culture flasks at 25°C. One group was fed (with fresh one-year-old pine twigs collected from healthy *P. massoniana* trees, with the needles removed) immediately after collection (“direct feeding”), whereas the second group was fed after a starvation period of 4 d (in a preliminary experiment, no nematodes departure from beetles within 4 days). In the present study, 41 *M. alternatus* were subjected to direct feeding and 37 were fed after a starvation period; the feeding of the beetles is shown in Fig. 1

### Extraction, counting and identification of PWN

The twigs of *P. massoniana* were replaced daily. The twigs had been fed by the beetles were cut into pieces and the PWN were extracted using the Bellman funnel method (Baermann, 1917). Twenty-four hours after extraction, the liquid at the bottom of the funnel was collected in a 10-mL centrifuge tube. The number of PWN was counted under a microscope (Zeiss Axio Lab. A1; Carl Zeiss, Göttingen, Germsny) under 10×4 field.

The molecular identification of nematodes was carried out to confirm that the extracted nematodes were PWN. 40 μL nematode liquid was transferred to a 1.5-mL Eppendorf tube with 40 μL of nematode lysis solution (Zhang et al., 2017) and 5 μL of protease K (Takara, Dalian, China). The mixture was then subjected to centrifugation at 2,000 ×g for 1 min before heating for 45 min at 65 °C in a constant temperature water bath, with the temperature then being raised to 95 °C for 10 min. Samples were then centrifuged at 12,000 ×g for 3 min, before collecting the supernatant and storing it at 4°C (Yang et al., 2020).

The 10-μL detection mixture consisted of 3.5 μL of the supernatant, 5 μL of Premix Ex Taq (Takara, Dalian, China), and 1.5 μL of a mixture of probe (5’-TGCAC GTTGT GACAG TCGT-3’) and primers (F: 5’-GAGCA GAAAC GCCGA CTT-3’, R: 5’-CGTAA AACAG ATGGT GCCTA-3’) in a PCR tube which was centrifuged at 2,000 ×g for 30 s. The detection mixture was analyzed by quantitative real-time PCR (qPCR) with the LineGene K Plus Real-Time PCR Detection System (FQD-48A; Hangzhou Bioer Technology Co. Ltd. Hangzhou, China) for the presence of PWN (constant temperature: 95 °C for 20 s; 40 cycles at 95 °C for 15 s, and 60 °C 129 for 20 s) (Yang et al., 2020).

### PWN motility and extraction percentage

PWN were extracted for a 4-h period from newly emerged *M. alternatus* and from beetles at 8 d after emergence, using the Bellman funnel method (Baermann, 1917). The PWN were washed with sterile water and treated in the dark for 10 minutes. Then, the motility of PWN was observed under a light microscope (Zeiss Axio lmager. M2; Carl Zeiss, Göttingen, Germsny) and videos were taken (Zeiss AxioCam HRc). The motility of the PWN carried by the beetles is shown in supplementary video A (the motility of the PWN carried by the newly emerged *M. alternatus)* and video B (the motility of the PWN carried by the beetles at 8 d after emergence).

PWN were extracted from ten newly emerged *M. alternatus* and from ten beetles at 8 d after emergence, with PWN being collected and counted every 6 h. The percentage of PWN for the total extraction was determined over different extraction periods (0—12 h, 12—24 h, 24—36 h).

### Transcriptome sequencing (Biomarker Technologies, Beijing, China)

#### RNA quantification and qualification

The total RNAs (three biological replicates of each of the two groups) of PWN were extracted, using the TRIzol Plus RNA Purification kit (Invitrogen, Waltham, MA, USA). RNA concentration was measured using NanoDrop^TM^ 2000 spectrophotometer (Thermo Fisher Scientific, Waltham, MA, USA). RNA integrity was assessed using the RNA Nano 6000 Assay Kit of the Agilent Bioanalyzer 2100 system (Agilent Technologies, CA, USA).

A total amount of 1 μg RNA per sample was used as input material for the RNA sample preparations. Sequencing libraries were generated using NEBNext^®^ Ultra™ RNA Library Prep Kit for Illumina^®^ (NEB, Ipswich, MA, USA) following manufacturer’s recommendations and index codes were added to attribute sequences to each sample.

#### Clustering and sequencing

The clustering of the index-coded samples was performed on a cBot Cluster Generation System using TruSeq PE Cluster Kit v4-cBot-HS (Illumia) according to the manufacturer’s instructions. After cluster generation, the library preparations were sequenced on an Illumina platform and paired-end reads were generated.

#### Quality control

Raw data (raw reads) of fastq format were firstly processed through in-house perl scripts. In this step, clean data (clean reads) were obtained by removing reads containing adapter, reads containing ploy-N and low quality reads from raw data. At the same time, Q20 (%), Q30 (%), GC-content (%) and sequence duplication level of the clean data were calculated. All the downstream analyses were based on clean data with high quality.

#### Comparative analysis

The adaptor sequences and low-quality sequence reads were removed from the data sets. Raw sequences were transformed into clean reads after data processing. These clean reads were then mapped to the reference genome sequence (https://www.ncbi.nlm.nih.gov/genome/?term=Bursaphelenchus+xylophilus). Only reads with a perfect match or one mismatch were further analyzed and annotated based on the reference genome. HISAT2 tools soft were used to map with reference genome.

#### Gene functional annotation

Gene function was annotated based on the following databases: Nr (NCBI non-redundant protein sequences); Nt (NCBI non-redundant nucleotide sequences); Pfam (Protein family); KOG/COG (Clusters of Orthologous Groups of proteins); Swiss-Prot (A manually annotated and reviewed protein sequence database); KO (KEGG Ortholog database); GO (Gene Ontology).

#### Differential expression analysis

Differential expression analysis of two conditions/groups was performed using the DEseq software. DEseq provide statistical routines for determining differential expression in digital gene expression data, using a model based on the negative binomial distribution. The resulting P values were adjusted using the Benjamini and Hochberg’s approach for controlling the false discovery rate (FDR). Genes with an adjusted *P*-value < 0.01 found by DEseq were assigned as differentially expressed.

#### KEGG pathway enrichment analysis

KEGG (http://www.genome.jp/kegg/) was used to identify biological pathways that were enriched in a gene list more than would be expected by chance. KOBAS (Mao et al., 2005) software was used to test the statistical enrichment of differentially expressed genes (DEGs) in KEGG pathways.

### Statistical analysis

Statistical analysis was performed using SPASS 19.0 (IBM, Armonk, NY, USA). One-way analysis of variance (ANOVA) was used to analyze the differences of the start departure time of PWN from *M. alternatus* feeding after starvation and direct feeding. Mann-Whitney test was performed to analysis the differences of difficulty level of extracted PWN from *M. alternatus* at 8 d after emergence and from newly emerged beetles.

## Results

### Effect of *M. alternatus* feeding behavior on PWN departure from beetle

After molecular identification, it was confirmed that the nematodes extracted in this study were all pine wood nematodes.

Data from the study into the time after *M. alternatus* emergence required for PWN start departure from the beetles are shown in Table 1. It can be seen that the mean (± standard deviaton, SD) start time for PWN departure from directly fed *M. alternatus* was 8.02 ± 1.96 d after beetle emergence, and there was no significant difference between male and female beetles fed directly (*P*=0.47). The mean start time for PWN departure from *M. alternatus* fed after starvation was 7.76 ± 2.28 d after beetle emergence, with no significant difference between male and female beetles (*P*=0.19). There was also no significant difference in the time that PWN started to departure from the beetles under the two feeding regimes (*P*=0.34) with no significant differences in the start time of PWN departure from female (*P*=0.58) or male (*P*=0.12) beetles under the two feeding regimes.

**Table 1.** The start date of pine wood nematode (PWN) departure from *Monochamus alternatus*

| Treatment | Direct feeding <sup>a</sup> |  | Feeding after starvation <sup>b</sup> |  |
| --- | --- | --- | --- | --- |
| Sex of beetles | ♀ | ♂ | ♀ | ♂ |
| Days after emergence of *Monochamus alternatus* | 5 | 6 | 5 | 5 |
|  | 6 | 6 | 6 | 6 |
|  | 6 | 6 | 6 | 6 |
|  | 6 | 6 | 6 | 6 |
|  | 7 | 6 | 6 | 6 |
|  | 7 | 6 | 6 | 6 |
|  | 7 | 7 | 6 | 6 |
|  | 7 | 7 | 7 | 6 |
|  | 7 | 7 | 7 | 7 |
|  | 7 | 8 | 7 | 7 |
|  | 7 | 9 | 7 | 7 |
|  | 7 | 9 | 8 | 8 |
|  | 7 | 9 | 8 | 9 |
|  | 8 | 10 | 9 | 9 |
|  | 8 | 10 | 9 | 9 |
|  | 9 | 10 | 9 | 12 |
|  | 9 | 11 | 10 |  |
|  | 9 | 12 | 11 |  |
|  | 10 | 12 | 12 |  |
|  | 10 |  | 13 |  |
|  | 10 |  | 14 |  |
|  | 13 |  |  |  |
| Average (mean ± SD) | 7.82±1.82 | 8.26±2.13 | 8.19±2.54 | 7.19±1.80 |
| | 8.02±1.96 | | 7.76±2.28 | |

a: The date of PWN start departure from *Monochamus alternatus* which was fed directly after emergence, with 41 beetles. b: The date of PWN start departure from *Monochamus alternatus* was which fed after a 40-d starvation period, with 36 beetles. There was no significant difference in the start date of PWN departure from beetles between the two treatments ( $P=0.34$ ), with no significant differences of PWN departure from female ( $P=0.58$ ) or male ( $P=0.12$ ) beetles.

The start date of PWN departure from *M. alternatus* fed directly was mainly concentrated in the period 6-10 d after emergence, with the 36 *M. alternatus* in this category accounting for 87.8% of the total. The start date of departure of PWN from *M. alternatus* fed after the 4–d starvation treatment was mainly concentrated in the period 6–9 d after emergence, with the 29 *M. alternatus* in this category accounting for 78.4% of the total. There was no significant difference between the two treatments in peak period of PWN start departure from beetles (*P* = 0.078).

These results indicated that the feeding behavior of *M. alternatus* had no significant effect on the departure of PWN. It also suggest that the consequent metabolic changes caused by different feeding treatments had no effect on the departure of PWN, and volatiles from pine twigs also had no effect on the departure of PWN (when the beetles were exposed to starvation, they were not stimulated by volatiles.).

### Comparisons of PWN motility

After dark treatment for 10 minutes (to simulate conditions in the body of the beetle), the PWN carried by the newly emerged beetles were less motile than the PWN which carried by the beetles at 8 d after emergence (supplementary videos A and B). At the same time, it was found that the PWN carried by *M. alternatus* at 8 d after emergence were easier to extract than the PWN carried by the newly emerged beetles (Table 2). A higher proportion of PWN being collected within 12 h from the beetles at 8 d after emergence than from the newly emerged beetles (*P* < 0.01). These findings suggest that the higher motility PWN were easier extracted from *M. alternatus* was associated with their departure from the vectors.

**Table 2.** The percentage of pine wood nematodes (PWN) collected in different extraction periods

|  | NO. | The time of PWN collected |  |  |
| --- | --- | --- | --- | --- |
|  |  | 0-12 h | 12-24 h | 24-36 h |
| Newly emerged beetles | 1 | 29% | 50% | 21% |
|  | 2 | 35% | 53% | 12% |
|  | 3 | 26% | 49% | 25% |
|  | 4 | 38% | 49% | 13% |
|  | 5 | 25% | 61% | 14% |
|  | 6 | 29% | 46% | 25% |
|  | 7 | 41% | 40% | 19% |
|  | 8 | 27% | 51% | 22% |
|  | 9 | 33% | 51% | 16% |
|  | 10 | 31% | 55% | 14% |
| Beetles at 8<br>day after | 11 | 80% | 15% | 5% |
|  | 12 | 84% | 11% | 5% |
|  | 13 | 91% | 6% | 3% |
|  | 14 | 81% | 13% | 6% |
|  | 15 | 77% | 16% | 7% |
|  | 16 | 75% | 17% | 9% |
|  | 17 | 72% | 20% | 8% |
|  | 18 | 81% | 15% | 4% |
|  | 19 | 83% | 12% | 5% |
|  | 20 | 70% | 23% | 7% |
| Mean ±SD | a | 31%±5.2% | 51%±6.0% | 18%±4.9% |
| | b | 79%±6.2% | 15%±4.8% | 6%±1.8% |

The mean ± SD percentage of PWN collected during different extraction periods relative to the total extraction number (a: extracted from newly emerged *M. alternatus*. b: extracted from *M.* *alternatus* at 8 d after emergence.). within 0-12 hours, the proportion of PWN extracted from beetles at 8 d after emergence was significant higher than that from newly emerged beetles ( $P <$ 0.01).

### Analysis of the PWN transcriptome from *M. alternatus* of different ages

Analysis of transcriptome sequencing results found that expression of the oxidative phosphorylation pathway genes of PWN, which extracted from their vector beetles at 8 d after emergence, was greatly up-regulated, compared with PWN which extracted newly emerged beetles (Fig. 2). A total of 36 differentially expressed genes were enriched in this pathway, 35 of which were up-regulated; this pathway was enriched with the most abundant up-regulated genes (Details: Supplementary material S1). This indicated that PWN carried by the beetles at 8 d after emergence were in an oxygen-rich environment. At the same time, the expression of genes associated with fatty acid degradation (fatty acid *β*-oxidation) in PWN extracted from the beetles at 8 d after emergence was also significantly up-regulated (**Fig. 3**) (Details: Supplementary material S2). The increase in fatty acid degradation and oxidative phosphorylation would generate a considerable amount of energy, which promote the enhancement of PWN motility.

**Fig. 2.**
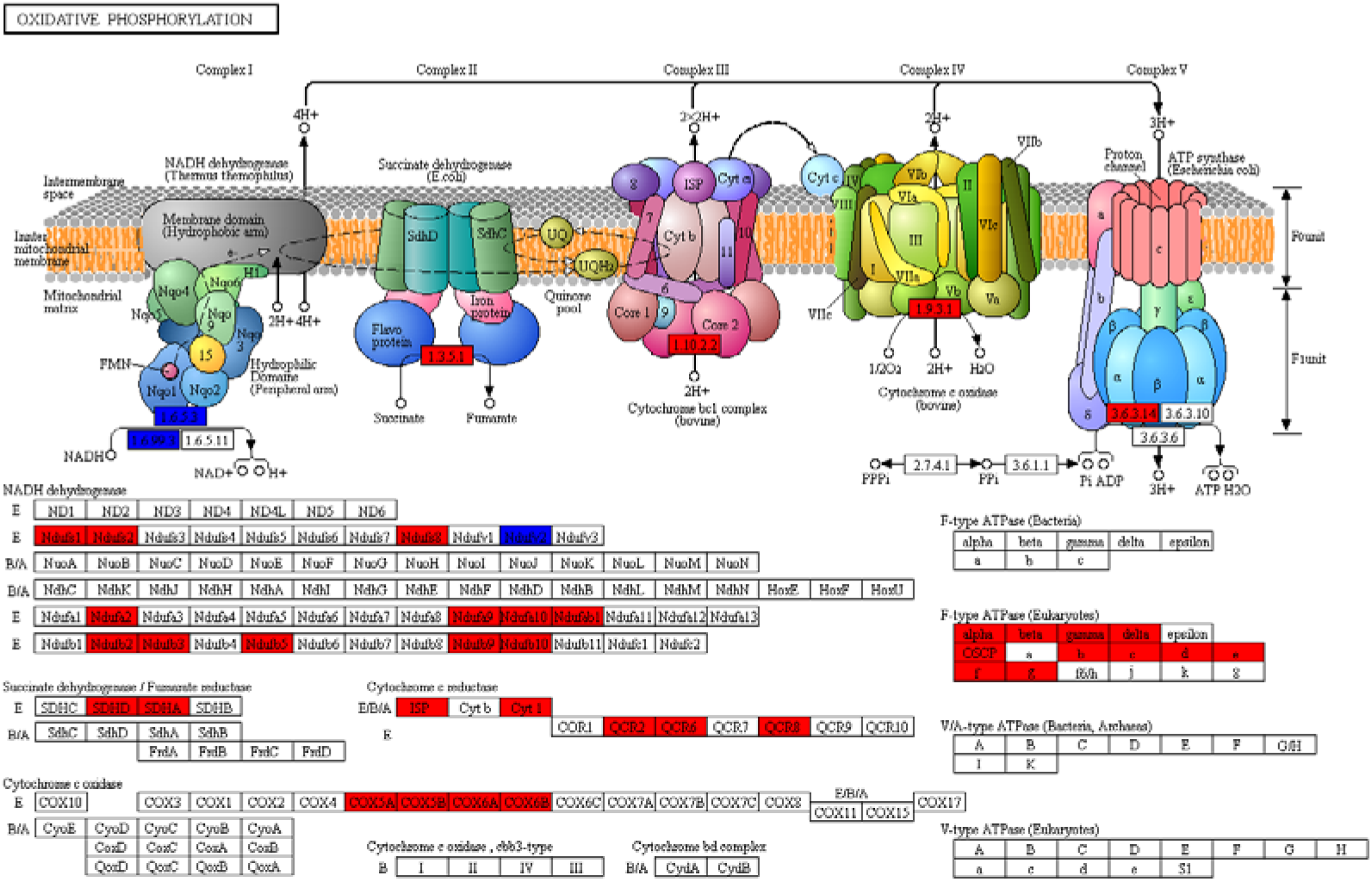
Oxidative phosphorylation pathway and gene expression profiles in pine wood nematodes, showing genes up-regulated or down-regulated in PWN extracted from 8-d beetles relative to newly emerged beetles. There were 36 differentially expressed genes (DEGs) in the oxidative pathway (in the analysis of protein subunits at the bottom half of the picture), the red block containing up-regulated genes (n=34), the blue block contains up-regulated genes and down-regulated genes. The two genes, encoding NADH hydrolase Ndufv2, was one down-regulated and another up-regulated (Details: Supplementary material S1).

**Fig. 3.**
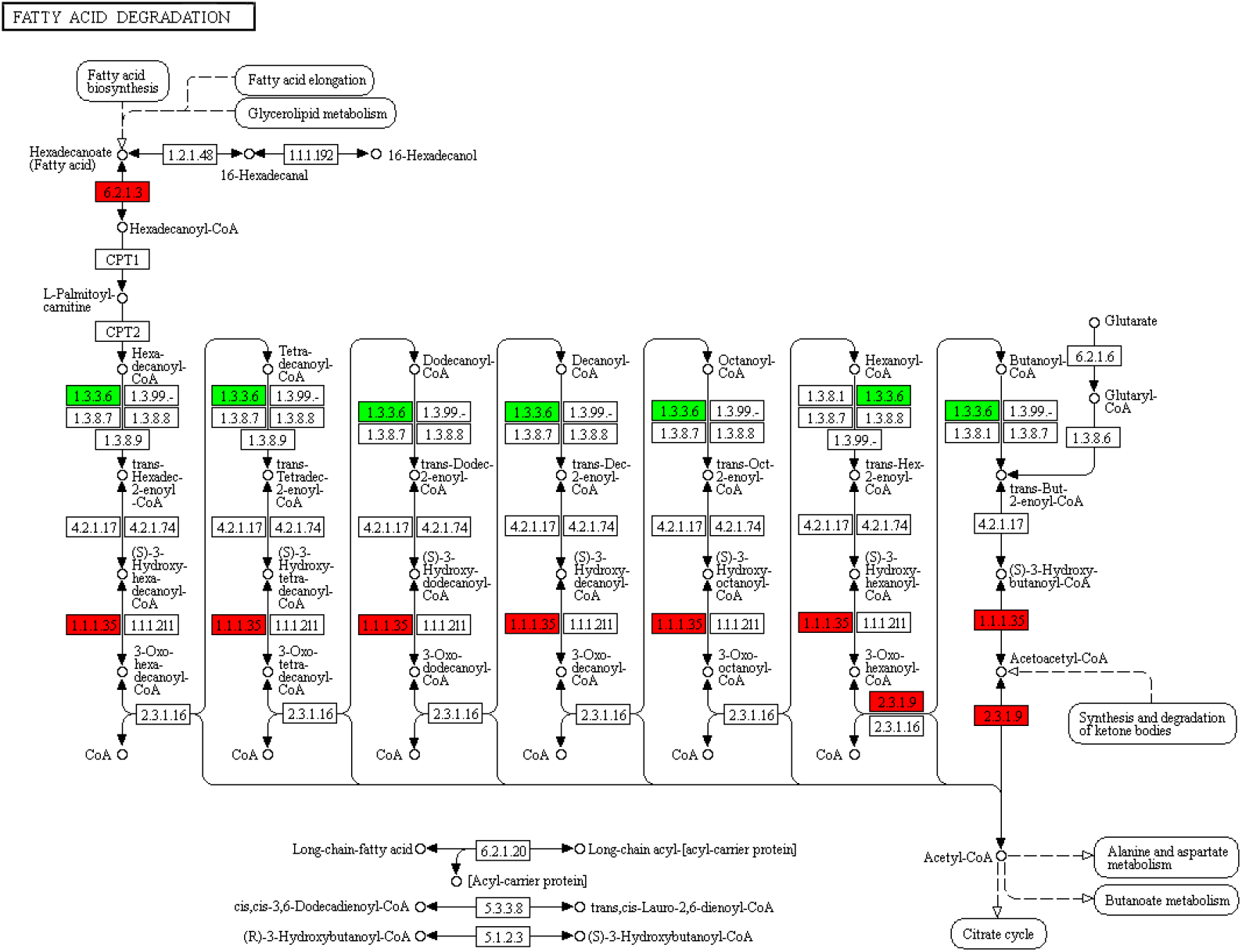
Fatty acid degradation pathway and gene expression profiles in pine wood nematodes, showing genes up-regulated (red) or down-regulated (blue) in PWN extracted from 8-d beetles relative to newly emerged beetle. The important genes (encoding long-chain acyl-CoA synthetase [EC:6.2.1.3], acetyl-CoA C-acetyltransferase [EC:2.3.1.9] and 3-hydroxyacyl-CoA dehydrogenase [EC:1.1.1.35]) for fatty acid *β*-oxidation were all up-regulated. Expression of the gene encoding acyl-CoA oxidase [EC:1.3.3.6], which inhibits fatty acid *β*-oxidation, was down-regulated.

## Discussion

A number of studies have investigated the factors which affect PWN departure from vectors, although differences in results have been reported. Stamps and Linit (1988) reported that neutral storage (NS) lipid was correlated with PWN exit and may contribute to PWN departure from vectors. A rolling fulcrum model of the integration of intrinsic (NS lipid) and extrinsic (volatiles) cues is proposed to explain the behavioral ontogeny of fourth-stage dispersive juveniles in relation to the beetle vector (Stamps and Linit, 2001). In the rolling fulcrum model, NS lipid content is proposed as a modifier of PWN response to beetle and tree produced volatiles, with juveniles with the lower NS lipid content being attracted to *β*-myrcene, a pine volatile, whereas PWN with the higher NS lipid content were attracted to toluene’ a beetle cuticular hydrocarbon (Stamps and Linit, 1998, 2001). Ishkawa et al. (1986) examined the attractant effect of volatile components in *Pinus densiflora,* to the PWN (dispersive PWN that extracted from *M. alternatus),* and assumed that *β*-mvrcene played an important role in the transmigration of the PWN from the vector to the pine tree and for the movement of the PWN inside the pine wood. However, in the present study, we believe that volatiles were not the cause of PWN departure from *M. alternatus.*

Provided autoclaved (without natural volatiles) and fresh pine twigs to *M. alternatus* respectively, the results suggested that PWN had a trait of spontaneous departure from *M. alternatus* and the volatiles from fresh *P. densflora* twigs repressed PWN departure from beetles (Aikawa and Togashi, 1998). Stamps and Linit (1988) examined the effects of various pine volatiles on PWN departure, suggested that PWN departure behavior possibly controlled by endogenous nature factors. Volatiles, previously thought to be attractants, may actually act more as aggregation stimulants than as attractants, with PWN moving randomly until the desired chemical is reached, whereupon movement decreases or stops (Futai, 1980; Stamps and Linit, 1998). Volatiles may only play a role in fourth-stage dispersal juveniles movement from beetles into pine wood (Stamps and Linit, 1988, 1998).

In the present study, feeding behavior of the beetles had no significant effect on the time of PWN departure from *M. alternatus.* The results also indicated that volatiles (starvation-treated beetles were not stimulated by volatiles) of pine and the metabolic changes of *M. alternatus* caused by feeding behavior had no effect on the departure date of PWN. The two main behaviors of *M. alternatus* following emergence are feeding and breathing. Have study reported that, after emergence, the respiratory rates of *M. alternatus* increased, which led to a rise in CO2 production rate (Wu et al., 2019). When the CO_2_ concentration in trachea reached a certain critical level, PWN began to escape from the beetle (Wu et al., 2019), with to avoid CO_2_ (Bretscher et al., 2008). In the present study, we observed that the PWN carried by the newly emerged beetles exhibited less motility than the PWN carried by the beetles at 8 d after emergence, with the PWN carried by *M. alternatus* at 8 d after emergence were easier to extract than the PWN carried by newly emerged beetles, suggest that the date of departure of PWN from vector might be related to their motility. PWN movement requires energy, lipid degradation provides the energy needed for exercise (Stamps and Linit, 1988,1998), with lipid fully degradation to produce large amounts of energy (*β*-oxidation) requiring sufficient O_2_. Respiratory behavior leads to a critical level of CO_2_ in the trachea of *M. alternatus* (Wu et al., 2019), with the O_2_ concentration in the beetle trachea also reaching a critical level when breathing (the concentration of O2 will increase when inhaling).

Tian et al. (2017) carried out analysis of metabolism-related genes and showed that the metabolic level of fourth-stage dispersive juveniles (carried by newly emerged beetles) was much lower than that in third-stage dispersive juveniles (in dead pine). This was due to the decreases in enzymatic activity in those pathways involving metabolism of glycolysis, oxidative phosphorylation, tricarboxylic acid cycle, gluconeogenesis, etc. (O’Riordan and Burnell, 1989). The expression of the sorbitol dehydrogenase gene in the fourth-stage dispersal juveniles (carried by newly emerged beetles) was up-regulated (Tian et al., 2017), this enzyme may be involved in ethanol fermentation under hypoxic conditions (Mcelwee et al., 2006), providing basic energy for PWN. These findings indicated that the PWN were in a hypoxic environment in the early stage after entering the trachea of *M. alternatus* (Tian et al., 2017). In the present study, the low motility of PWN carried by newly emerged beetles also supported that PWN are in a state of low oxygen and low energy.

In the present study, the greater motility of PWN (carried by *M. alternatus* at 8 d after emergence) were easier departure from *M. alternatus.* The greater motility showed that, under these circumstances, PWN had a strong metabolism and abundant energy. A large amount of lipid degradation in PWN, after a period of emergence of the beetle (Stamps and Linit, 1988), and oxidative phosphorylation requires sufficient oxygen sustained stimulation. In the present study, transcriptome sequencing found that expression of the genes associated with oxidative phosphorylation level of PWN carried by the beetles at 8 d after emergence was significantly higher than that in PWN carried by newly emerged beetles. The thermodynamic power of the oxidative phosphorylation electron transfer chain comes from the strong affinity of oxygen for electrons (Wang et al., 2002), indicating that PWN in the trachea of beetles (at 8 d after emergence) obtained sufficient oxygen, with the oxygen obtained by PWN can only come from the respiration of the beetles.

In summary, our view is that the increased of respiration after adult vector emergence (Wu et al., 2019) causes an increase in O_2_ concentration in the trachea, which causes the increase of oxidative phosphorylation level in PWNAn effect of pine volatiles on departur, An effect of pine volatiles on departuree which, in turn, then causes the degradation of lipid (oxidative phosphorylation and *β*-oxidation are coupled (Wang et al., 2002)). This increase in oxidative phosphorylation in PWN releases energy, which increases the motility of PWN, and is the direct internal cause of their departure from the vector beetle, with the increased motility being the factor necessary for PWN departure from the vector.

## Acknowledgement

This study was supported by the Funding information the National Key R&D Program of China, Grant/Award Number: 2017YFD0600104.

