## Supplementary figures and images for "The effect of feeding behavior of *Monochamus alternatus* (Coleoptera: Cerambycidae) on the departure of pine wood nematode, *Bursaphelenchus xylophilus* (Nematoda: Aphelenchoididae)"

### ko00071.png

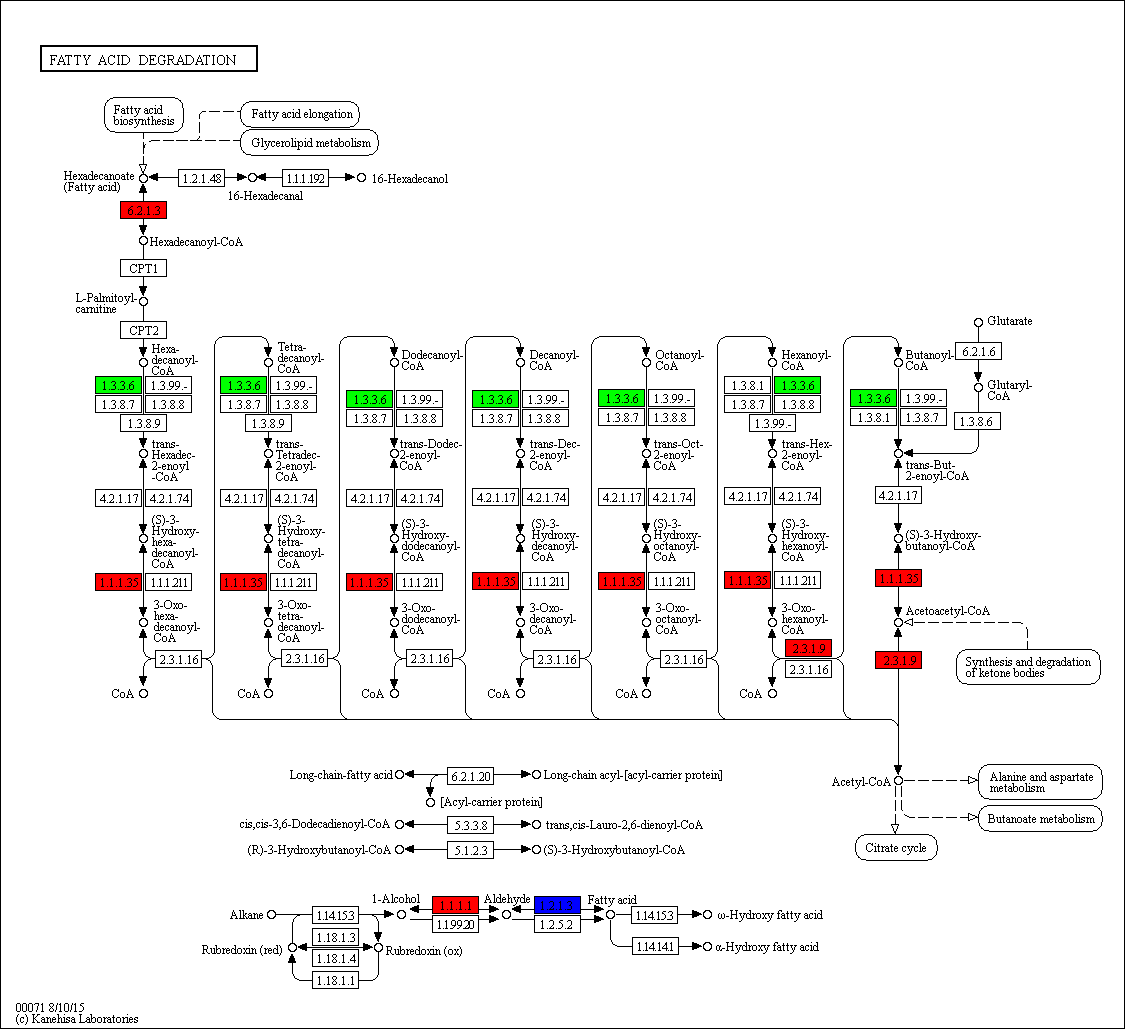

### ko00190.png

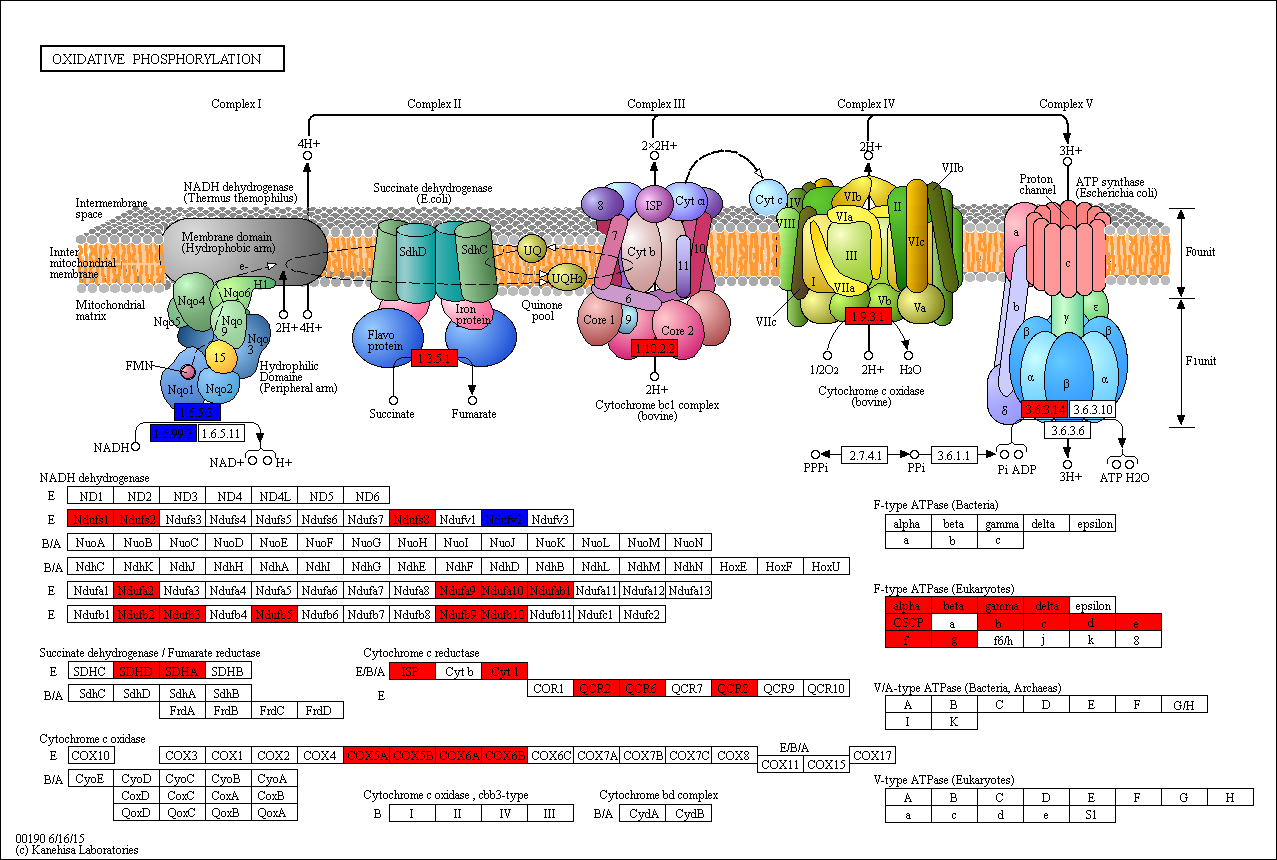
